# *Daphnia s*tressor database: Taking advantage of a decade of *Daphnia* ‘-omics’ data for gene annotation

**DOI:** 10.1101/444190

**Authors:** Suda Parimala Ravindran, Jennifer Lüneburg, Lisa Gottschlich, Verena Tams, Mathilde Cordellier

## Abstract

Gene expression patterns help to measure and characterize the effect of environmental perturbations at the cellular and organism-level. Complicating interpretation is the presence of uncharacterized or “hypothetical” gene functions for a large percentage of genomes. This is particularly evident in *Daphnia* genomes, which contains many regions coding for “hypothetical proteins” and are significantly divergent from many of the available arthropod model species, but might be ecologically important. In the present study, we developed a gene expression database, the *Daphnia* stressor database (http://www.daphnia-stressordb.uni-hamburg.de/dsdbstart.php), built from 90 published studies on *Daphnia* gene expression. Using a comparative genomics approach, we used the database to annotate *D. galeata* transcripts. The extensive body of literature available for *Daphnia* species allowed to associate stressors with gene expression patterns. We believe that our stressor based annotation strategy allows for better understanding and interpretation of the functional role of the understudied hypothetical or uncharacterized *Daphnia* genes, thereby increasing our understanding of *Daphnia*’s genetic and phenotypic variability.

## Introduction

Environmental health is an important determinant of human health^1^ and natural habitats are under increasing threat from human activity. Industry and private households release chemicals in the environment, and human activity causes habitat fragmentation and warming, thus affecting ecosystems in their entirety. A wide diversity of organisms undergo changes in behavior, morphology and physiology by responding to biotic and abiotic factors in their environments^2–5^.

When an organism is subjected to one or multiple environmental perturbations, its gene expression and metabolic profiles are altered. Plastic alterations of the physiology, behavior or morphology are observed. Gene expression patterns provide clues about the biochemical pathways that are affected by a specific stressor^6^. Organisms under stress may show unique signatures of gene expression patterns associated with a stress response as seen in plants^7^ and rat liver^8^, for example. These unique patterns of gene expression can be used as indicators of specific environmental stressors.

Quantifying and comparing the gene expression profiles before and after exposure to a stressor is the most widely used approach to understand the physiological consequences of stressors. The central goal of most gen-/transcript-omic experiments is to obtain sets of differentially expressed genes and interpret the observed patterns. Complicating interpretation is uncharacterized gene function for a large percentage of genomes. This is particularly evident in crustaceans which are biologically diverse organisms and have been the subject of investigation for hundreds of years because of their important role in ecology, evolution and ecotoxicogenomics^9^. Amongst crustaceans, gene prediction tools in *Daphnia* genomes identified many coding regions but they are substantially divergent from the available genomes of arthropod model species, and annotation beyond “hypothetical protein” was not possible. Although *Daphnia*’s ecology is intensively studied, still little is known about the molecular basis of responses to environmental stressors.

*Daphnia* is an ecologically important model organism distributed throughout the world in a variety of habitats ranging from lakes to ponds and from hypersaline to freshwater systems. They are primary grazers of algae, bacteria and protozoans and primary forage for fish^10^ as well as larger invertebrates. As a consequence, *Daphnia* are considered to be the sentinel species of freshwater environments where their decline is proportional to the presence of environmental perturbations^11^. Several studies^12–15^ in *Daphnia* exist which quantify their gene expression profiles in response to stressors affecting their phenotype (i.e. behavior, metabolism, life history traits and morphology). These observations have become more feasible with the advent of *Daphnia* genomics, which have seen rapid advancements in the last decade with the availability of the first genomes of *D.pulex^16^, D.magna, D.similoides* and transcriptome of *D. galeata^17^.* The wFLEABASE^18^ developed by the *Daphnia* Genomics Consortium (DGC) currently serves as a hub for obtaining gen-/transcript-omic *Daphnia* data. However, a *Daphnia* repository dealing with experimentally validated gene expression and its response to environmental stressors is lacking.

By taking advantage of comparative genomics approaches, in the present study we wanted to identify transcript-specific stressors in *D. galeata* using the functional annotation data available from a previous study^17^. We observed that most of the retrieved *D. galeata* transcripts annotations pointed to “hypothetical proteins” predicted in *D. pulex* or the function was “unknown”. To overcome this challenge, we identified studies dealing with *Daphnia* gene expression using three different search strategies. We manually mined the data to identify differentially expressed genes and transcripts for each stressor. Using a homology approach, we were able to annotate *D. galeata* transcripts to *Daphnia* specific genes in literature known to respond to one or more stressors through regulatory changes. We further identified transcripts/genes that were responding to many stressors thereby forming a general stress response and also identified stress-specific transcript/genes.

The results of our literature review were implemented into a database called *Daphnia* stressor database (http://www.daphnia-stressordb.uni-hamburg.de/dsdbstart.php). To the best of our knowledge, this study is a first attempt to integrate the wealth of information readily available by using a comparative genomics approach. We believe this long overdue resource will be of use to researchers working in the fields of ecotoxicology, environmental health, evolutionary biology and environmental genomics, and serve as a basis for further investigations in *Daphnia* research.

## Results and Discussion

### Number of studies obtained from each search strategy

A total of 676 studies were obtained from all the three search strategies as described (in methods section). Of them, 47 studies came from FunctionSearch method, 74 studies came from Mendeley search and 555 came from automatic retrieval using twitter. We excluded studies not dealing with gene expression in *Daphnia* and studies from which information was not retrievable (due to language restrictions, intellectual property rights, primer sequences and/or gene names instead of IDs were mentioned). After screening the literature database with the above-mentioned criteria and checking for duplicates, gene expression data specific to *Daphnia* was retrievable from 90 studies (Supplementary Table S1) and used for the present meta-analysis (Figure 1a).

**Figure 1:**
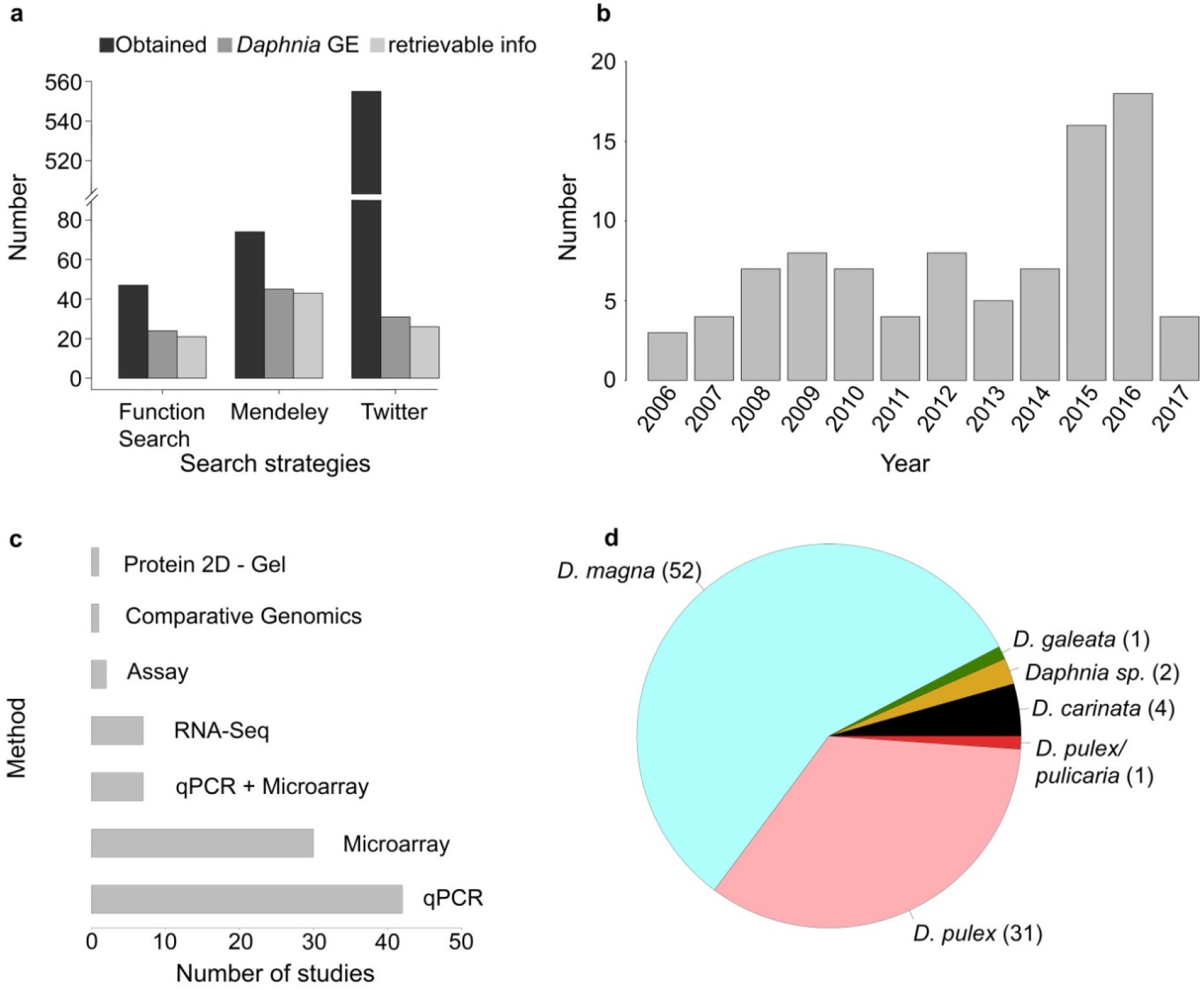
Literature statistics for meta-analysis. (a) Barplot indicating the number of studies obtained for each search strategy used in this meta-analysis. Bars in black represent all literature obtained in each search strategy. Bars in dark grey represent the number of studies specific to *Daphnia* and gene expression; bars in light grey represent the number of studies from which data was retrievable for analysis. (b) Timeline for studies on gene expression in *Daphnia*. (c) Barplot showing the number of studies available for different analysis method used in the literature. (d) Pie chart showing the number of studies available for each *Daphnia* species.

### Timeline of literature, species information and methods used

After limiting the literature database to gene expression studies in *Daphnia*, we obtained literature from the year 2006 until mid- 2017 (Figure 1b). About 43 studies used qPCR, thus limiting their analysis to only a few genes (Figure 1c). After the advent and further development of DNA microarray at the dawn of the millennium, researchers took advantage of the technology to perform large-scale gene expression studies. In our survey, 30 studies used microarray technology to analyze gene expression patterns in *Daphnia*. Seven studies combined both microarray and qPCR approaches. The advent of high-throughput sequencing (HTS) techniques and affordable sequencing costs made *Daphnia* genomes and transcriptomes a reality. In this literature survey, we found 7 studies using RNA-seq to analyze whole-transcriptome expression profiles. Very few of the studies (3 studies) in the literature database used comparative genomics approaches, biochemical assays and 2D- gel electrophoresis techniques.

The majority of the studies (52 out of 90) (Figure 1d) dealt with *D. magna*, followed by *D. pulex* (31 out of 90) and very few on other *Daphnia* species like *D. carinata* (4 studies), *Daphnia sp.* (reviews, 2 studies), *D. galeata* and *D. pulicaria* (one study each).

### Daphnia genes associated to a stressor in literature

After retrieving the gene IDs regulated in response to a stressor from literature, ~21% of the 30939 genes^16^ with a DAPPUDRAFT id (e.g.: DAPPUDRAFT_xxxxxx) present in *D. pulex* were associated to a stressor, i.e. were differentially expressed under this stressor (Figure 2a). Among them, the expression of ~84% of the genes was affected by one stressor, 13% of the genes by two stressors, 2% by three stressors and less than 0.5% between 3 and 9 stressors.

**Figure 2:**
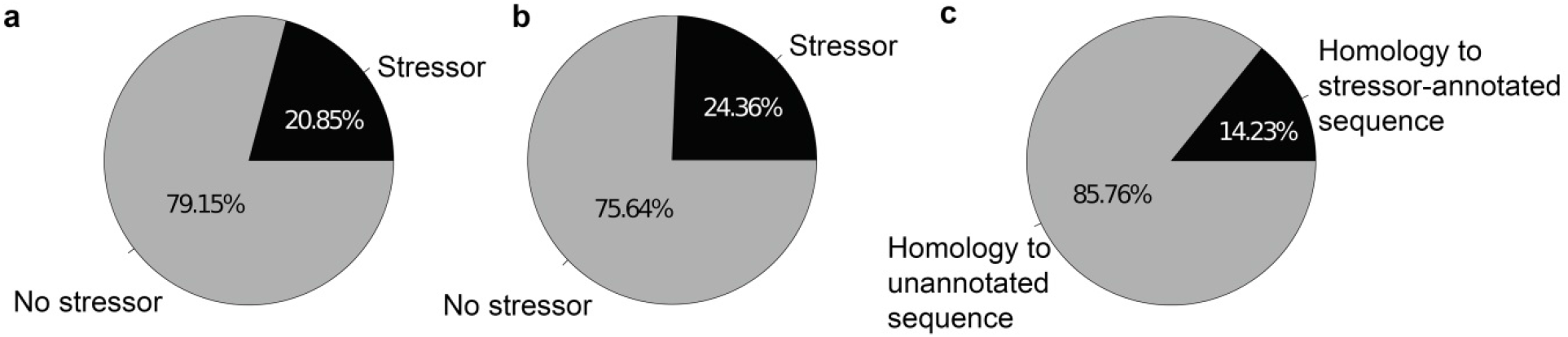
Pie charts showing the percentage of genes associated to a stressor in the literature database for (a) *D. pulex* (with DAPPUDRAFT nomenclature IDs); (b) *D. magna* (with Dapmaxxxxxxxx nomenclature IDs); (c) *D. galeata* (test case used in the present analysis).

For *D. magna*, ~25% of the 29127 genes were associated to a stressor that came from microarray studies (Figure 2b). Among them, ~51% of the *D. magna* genes associated to a stressor were responding to one stressor, ~23% to two stressors, ~9% to three stressors and less than 5% of genes were responding to between 4 and 31 stressors.

To represent the shared response between stressors, we used a circularized plotting using circlize package^19^ in R. Every stressor field represents the list of genes shown to be differentially expressed. A line linking one stressor field to another implies that a gene is differently expressed for both stressors. The link density thus makes the non-specificity of some regulatory responses evident. For *D. magna*, stressor fields were often highly linked to other stressors (Figure 3). This leads to the conclusion that the genes might be showing a general stress response. In *D. pulex* (Figure 4), some stressors showed very few links (i.e. temperature, light: dark cycle, phosphorous). In this case, the stressor response might be specific and even used for environmental diagnosis.

**Figure 3:**
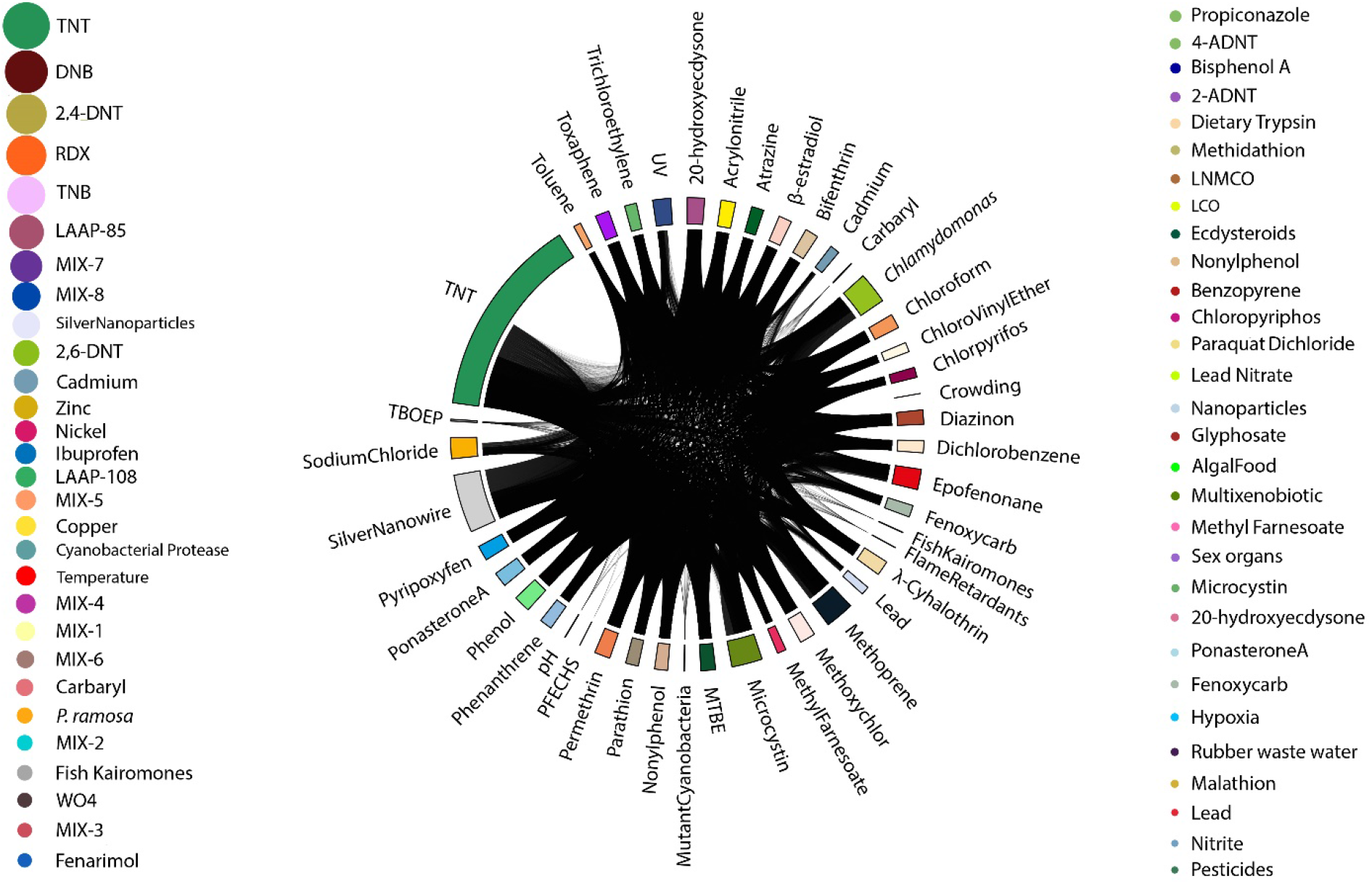
Circular and bubble plot showing the number of genes associated to one or more stressors in *D. magna*. Every stressor field on the circular plot represents the list of genes identified to be differentially expressed, and its size is proportional to the number of genes. A line linking a field to another one means that a gene is differently expressed for the other stressor as well. For genes that followed other gene ID nomenclature for *D. magna*, stressors are represented through bubbles on either side of the circular plot, and the bubble size is proportional to the number of genes.

**Figure 4:**
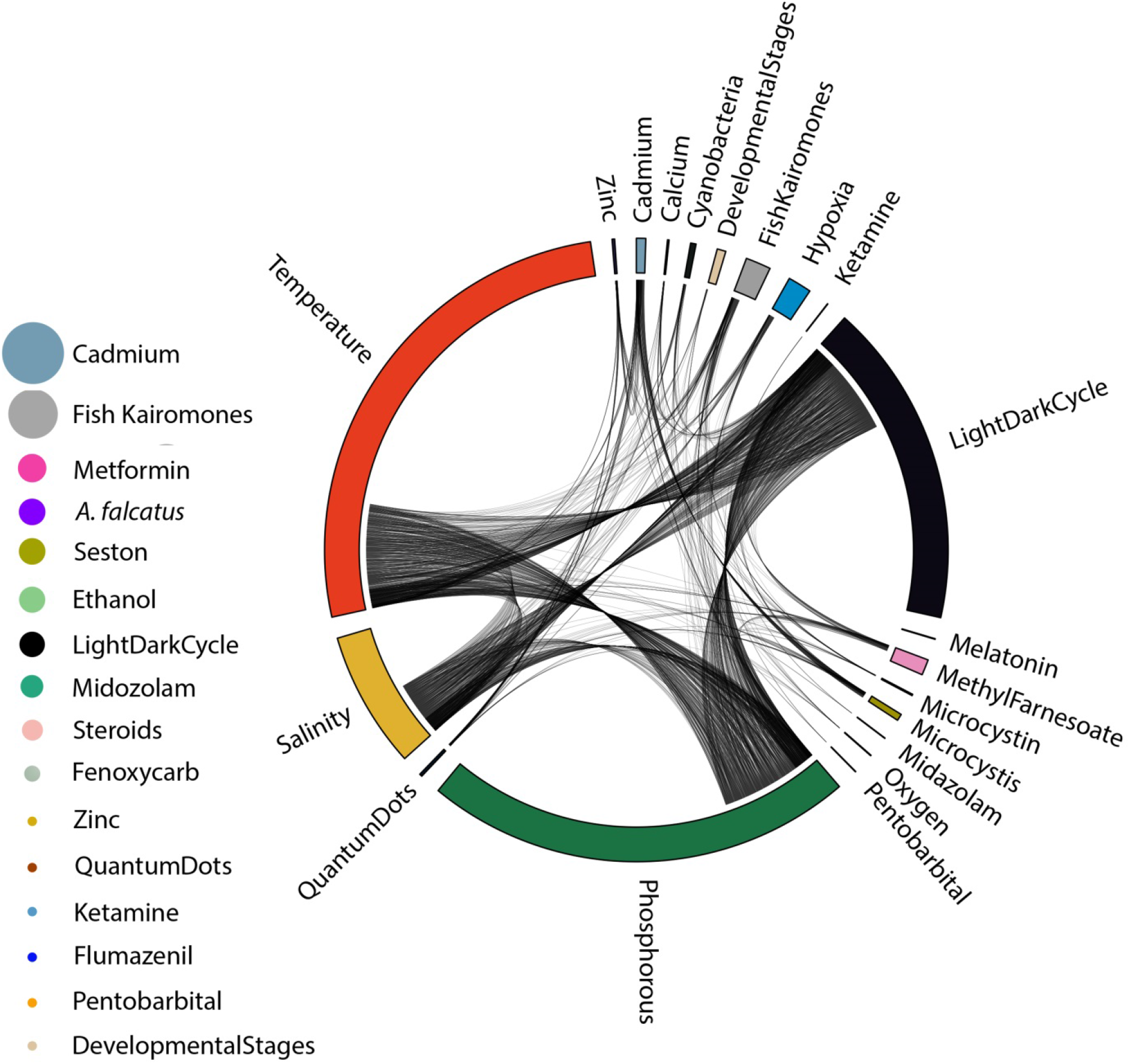
Circular and bubble plot showing the number of genes associated to one or more stressors in *D. pulex*. Every stressor field on the circular plot represents the list of genes identified to be differentially expressed, and its size is proportional to the number of genes. A line linking this field to another one means that a gene is differently expressed for this stressor as well. For genes that followed other gene ID nomenclature for *D. pulex*, stressors are represented through bubbles on the left side of the circular plot, and the bubble size is proportional to the number of genes.

A number of studies did not use the “DAPPUDRAFT” or “Dapmaxxxxxxxxxxx” nomenclature, yet they were dealing with stressors associated to *D. pulex* and *D. magna,* respectively. These genes and their associated stressors were visualized using bubbles (Figures 3, 4), where each bubble corresponds to one stressor and their size is proportional to the number of genes associated to that stressor.

### D. galeata genes associated to a stressor using homology approach

The assembly of *D. galeata* was used as a test case and gives an idea of the power of our approach for newly sequenced and assembled *Daphnia* or cladoceran/crustacean genomes. After performing BLAST analysis for each of the 32903 *D. galeata* transcripts, we observed significant hits (eval < 0; identity ≥ 50%) for 4684 transcripts (Figure 2c). Among them, 4478 *D. galeata* transcripts shared homology with nucleotide sequences, 61 transcripts with protein sequences in the database and 145 transcripts had both a nucleotide and protein sequence hit from the database (Figure 5a).

**Figure 5:**
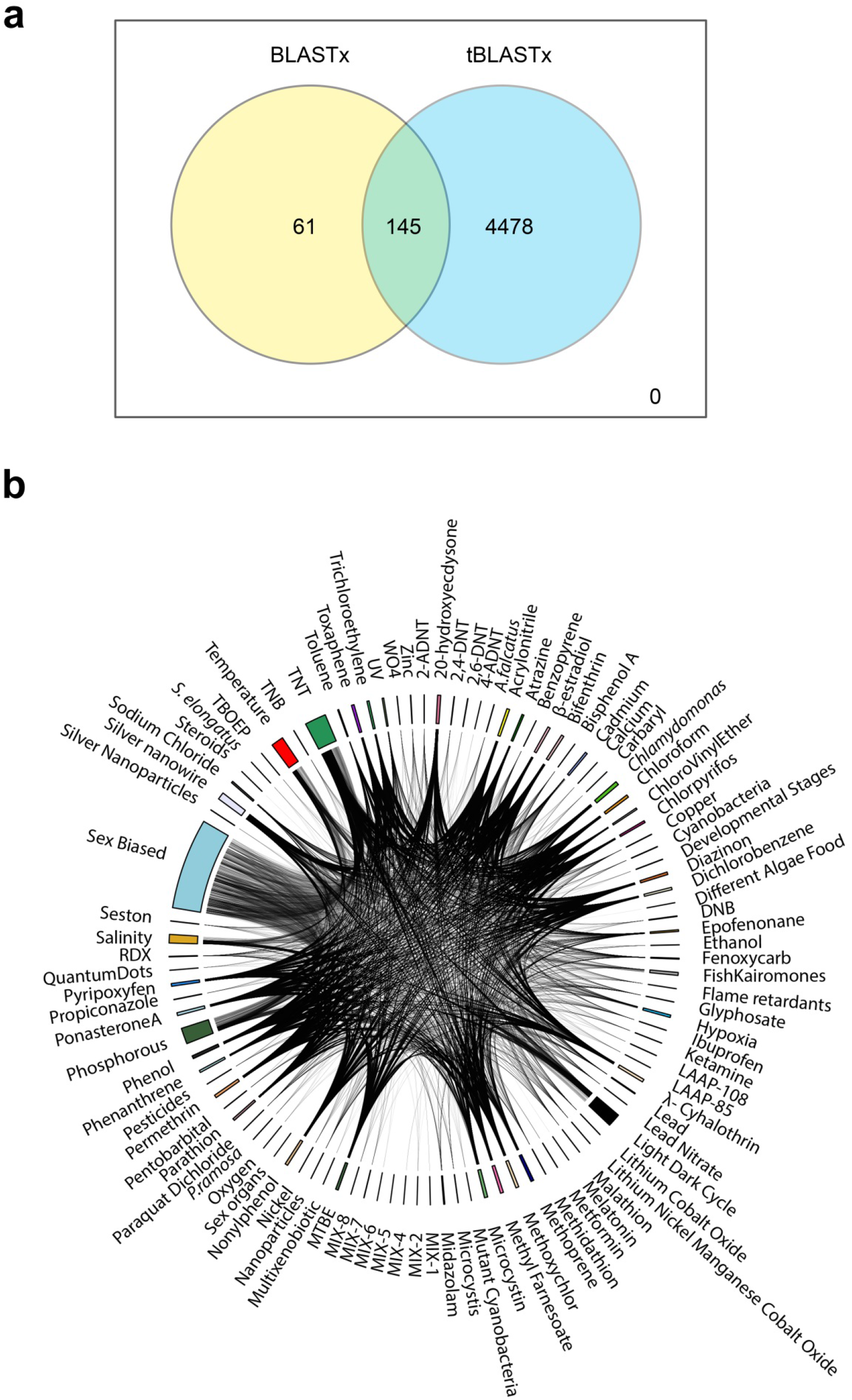
(a) Venn diagram representing the overlap between blastx and tblastx analysis used for annotating *D. galeata* transcripts. (b) Circular plot showing the *D. galeata* transcripts presenting sequence homology to at least one sequence in the Daphnia stressor db, classified according to the stressor affecting their expression. Every stressor field represents the list of genes identified to be differentially expressed. A line linking this field to another one means that a gene is differently expressed for this stressor as well.

For *D. galeata*, ~64% of the transcripts matched a gene responding to a single stressor (as revealed by our comparative approach), ~20% a gene responding to two stressors, ~6% a gene responding to three stressors. Less than 4% were matched to genes associated to between 4 and 31 stressors (Figure 5b).

Although our approach is not *per se* an experimental validation of the associations between stressor and gene expression, it allows to further our knowledge and to formulate hypotheses. Within the *D. galeata* transcripts, a substantial amount were homologous to *D. pulex* genes. However, these were themselves only annotated with a “hypothetical function”. Our data mining approach allowed for their environmental annotation, which in turn provides an interpretation help as to the role of 4159 *D. galeata* transcripts homologous to these sequences.

### Searching the database

Database users should be able to retrieve information on experimentally validated differentially expressed genes in *Daphnia*. A search tool is provided for searching based on the following fields (1) gene ID differentially expressed in reponse to a stressor (2) stressor (3) title of the published research paper (4) authors of the study (5) technique used (6) study species (7) publication year and sequence of the respective gene. The user can query the database using a single field or a combination of multiple fields. The “advanced search” provides a refined way to search the database using several combinations of keywords using logical operators “+” / “−”, which represents “AND” / “OR”, respectively.

The result table includes a listing of all genes matching the search terms, and is interactive: hyperlinks behind the gene ID and the articles title lead to the sequence and the PubMed page of the publication, respectively.

### BLAST

To enable searches based on sequence homology and thus extend the use to studies using *de novo* assemblies or array probes without typical geneIDs, we implemented a BLAST tool in the database. This tool can be used for similarity-based search of any query sequences, protein and nucleotide. It is to be noted, though, that the database contains only genes responding to stressor and is therefore not an alternative to repositories such as wFLEAbase. The user can submit one or more sequence(s) in fasta format in the search field and set up thresholds for the e-value, among others.

### Limitations of the study and recommendations to the community

Although an extensive literature search for retrieving the differentially expressed *Daphnia* genes in response to a stressor was performed, we might have missed studies that did not turn up using the three search strategies followed. However, since the *Daphnia* stressor database is publicly available, data will be added manually and updated on a regular basis. As more data gets published or where our search strategy failed to retrieve the literature, we intend to include them in the database for the benefit of the *Daphnia* research community. Researchers also can contact the authors to include their experimentally validated gene expression data.

Because our study did not include a re-analysis of all the datasets, we chose to take the inferred changes in gene expression at face value. The caveat is that while most cited studies were following standard practice at the time of the publication i.e. using state of the art statistical tools and setting similar thresholds for significance, some didn’t apply multiple testing corrections. In total, 8 of the 43 studies based on either microarray or RNA-seq procedures did not use multiple testing corrections, and are identified accordingly in our database. The results aggregated in our study should therefore be reviewed critically when used to formulate hypotheses, and re-analyzed if necessary. Eventually, and once highly contiguous reference genomes are available for these species, the data shall be re analyzed. This will be possible in the near future, as two genomes for *Daphnia pulex* are readily available^20,21^ two genomes for *Daphnia magna* as well^22,23^, and the *Daphnia galeata* genome is being assembled as we speak.

Gene expression in *Daphnia* has been extensively studied in the last decade and the present study brought a few weaknesses and strengths to light. *Daphnia* researchers can rely on very extensive data compared to researcher communities focusing on other taxa, especially regarding ecologically relevant stressors. The high numbers of stressors and studies constitute a wealth of information and are an invaluable resource for the future. However, the data mining procedure highlighted a few shortcomings, some of which can be easily remedied. Our recommendations for present and future publications are contained in three words: traceable, searchable, and sustainable. First, publications should use traceable IDs to allow for cross referencing and good search results based on gene IDs. Whenever possible, gene IDs already in use in long term online repositories such as GenBank or Uniprot IDs should be used. Several studies that used qPCR-based approaches provided result tables with gene names specific to the paper and not in use otherwise, , meaning they were *de facto* non-traceable and no matching sequence could be found. Therefore, using already existing gene IDs (instead of gene names) would be tremendously helpful to include these results in the database and influence their citation record. Second, the results should be searchable, and preference should be given to tables in a universal readable format (csv, tsv, tab delimited txt, spreadsheets to some extent), instead of heatmaps saved as a graphic. Last but not least, sustainability is an issue affecting all publications and becomes increasingly important with the high volume of data produced, associated with publication on one side, and high personal turnover inherent to research on the other side. Both factors are challenging, but solutions already exist in the form of above mentioned repositories. Such centralized repositories, the Expression Atlas^24^ and GEO Profiles ^25,26^, gather gene expression data but only for a few selected species. Following these recommendations would allow a better integration of new and older results across laboratories working with hypothesis, and increase the visibility of the *Daphnia* research community.

## Conclusion

Stress response is crucial for an organism’s survival in natural habitats. Identifying and functionally annotating ecologically relevant genes is an important aspect of ecotoxicological and evolutionary studies. Signature patterns of gene expression and their responses can be used as markers of environmental stressors. Traditional approaches often subject the genes to laboratory induced stressors and quantify their expression patterns. In the present study, we took advantage of comparative genomics approaches to identify transcript specific stressors in *D. galeata* using data mined from experimentally validated gene expression studies in *Daphnia*. The present database on gene expression studies in *Daphnia* will be useful for the community in designing stressor specific experiments and evaluating genes in response to environmental perturbations. Approaches such as the overrepresentation analysis presented by Herrmann *et al* ^27^ and Bowman *et al* ^28^ can be extended to other stressors than temperature. Last, our database will also help interpreting the results of studies on adaptation in natural populations as well as ecological experiments.

## Methods

### Identification and selection of eligible Daphnia-specific gene expression datasets for this study

We used three different strategies to retrieve literature that analyzed gene expression data in *Daphnia*.

#### a. Literature search based on functional annotation

Using the functional annotation data obtained from a previous study^17^ on *D. galeata*, we searched for literature using the keywords “*Daphnia*” + “stress” along with the predicted function for each transcript.

#### b. Literature search based on keywords in Mendeley

We used keywords “*Daphnia*” + “Gene Expression” in Mendeley reference manager and obtained sets of *Daphnia* specific gene expression papers.

#### c. Twitter based search

We automatically retrieved all literature posted in @wtrflea_papers twitter handle using a modified python script (available in our github repository), and subsequently retained only studies about gene expression.

A consolidated literature list was created with studies obtained from all three approaches mentioned above. Studies were excluded from the analysis for the following reasons: different language than English, no traceable gene names, or only primer sequences instead of gene IDs. The following information was extracted from each identified study in the consolidated list: (i) gene IDs that are differentially expressed in response to a stressor or condition; (ii) method used for the analysis; (iii) stressor(s) used in the study; (iv) study species.

### Sequence retrieval

Each retrieved gene or protein ID identifier was manually checked to see if they followed the NCBI (e.g. AB021136), Uniprot (e.g. E9FXA0), or wFLEABASE (e.g. DAPPUDRAFT_224348) nomenclature. Sequences for all obtained gene IDs were then retrieved from the respective databases (NCBI / Uniprot / wFLEABASE) using the trial version of iMacros tool (https://imacros.net/overview/), which is an automated tool for web scraping and data extraction. For gene IDs that followed a *D. magna* ID nomenclature (e.g.: Dapma7bEVm000787t1), sequences were directly obtained from the CDS file available on wFLEABASE (http://server7.wfleabase.org/genome/Daphnia_magna/openaccess/genes/Genes/earlyaccess/dmagset7finloc9b.puban.cds.gz).

For microarray-based studies on *D. magna*, the array accession number (eg: A-GEOD-16518) cited in the publication was queried in Gene Expression Omnibus^25^ (GEO). Reporter sequences were retrieved and a local BLAST was performed against the *D. magna* CDS sequences mentioned above. Hits that had a similarity ≥ 90% were considered for the next steps in the analysis.

All retrieved sequences were manually sorted into a “protein” or “nucleotide” categories using a perl script available in our Github repository.

### Local sequence database creation and BLAST analysis

Two fasta files were created, one with all nucleotide sequences and one with all protein sequences. Redundant sequences from the two fasta files were excluded using the biopython script Sequence Cleaner (https://biopython.org/wiki/Sequence_Cleaner) with the default parameters. Thus, two non-redundant local databases were created with 28259 nucleotide sequences and 102 protein sequences for BLAST^29^ analysis.

Each of the 32903 *D. galeata* transcripts was used to query the local non-redundant nucleotide (tblastx) and protein (blastx) databases created with the gene IDs obtained from literature. Hits with evalue (eval) ≤ 0 and identity percentage ≥ 50% were considered and the stressors corresponding to their respective subject ID were assigned to the *D. galeata* transcripts.

A general workflow of the entire meta-analyses is represented in Figure 6.

**Figure 6:**
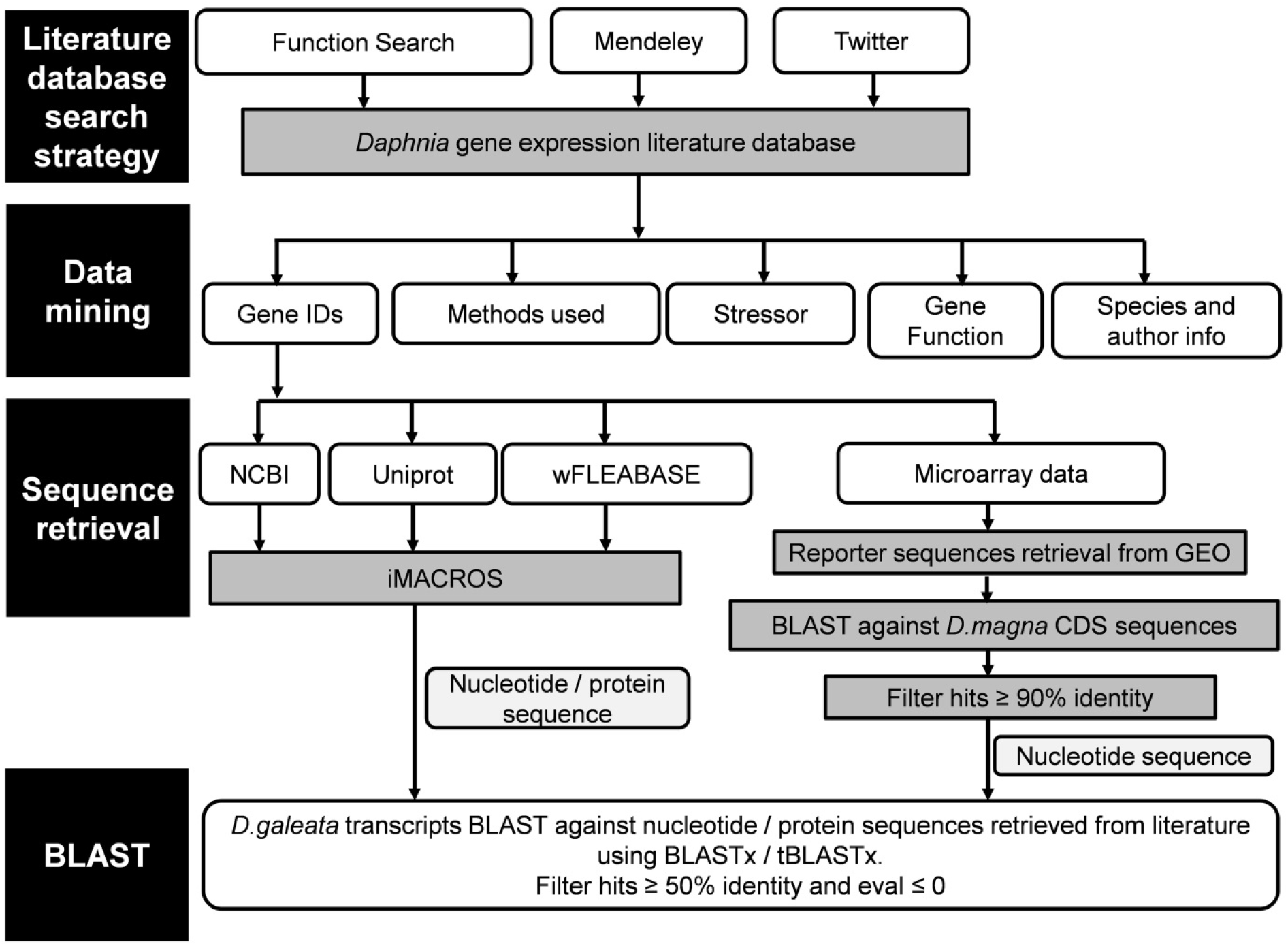
Workflow of Daphnia-specific gene expression followed in the present study.

### Database web interface

*Daphnia* stressor database (http://www.daphnia-stressordb.uni-hamburg.de/dsdbstart.php) is built on Mariadb (10.0.31-MariaDB) at the backend whereas the frontend is built using HTML, PHP and CSS3. To allow users to identify gene-specific stressors in *Daphnia* using a homology approach, we implemented a BLAST function on the database using SequenceServer^30^.

### Statistical analysis

All statistical analyses were performed using custom scripts in R^31^. These scripts are available in our Github repository PopGenHamburg/DaphniaStressordb.

## Supporting information

List of gene expression studies in Daphnia

## Supporting information

## Supporting information File 1

Table S1: List of gene expression studies in *Daphnia*, authors of the paper, year of publication, stressors used, species, experimental technique used in the study.

Custom scripts available at Github in the repository PopGenHamburg/DaphniaStressordb

## Acknowledgements

We would like to thank the Volkswagen Stiftung (grant to MC) for financial support. All participants to the 11^th^ Symposium on Cladocera 2017 played a role in shaping the current database. S. Thiemann, M. Herrmann, Y. Lu, J. Asselman and M. Möst provided useful comments on earlier versions of the database and the Rechenzentrum of Universität Hamburg provided assistance for developing the database. L. Orsini and two anonymous reviewers provided feedback on an earlier version of this manuscript. This work benefits from and contributes to the *Daphnia* Genomics Consortium.

## Author Contributions

SPR and MC designed the study and reviewed the literature. SPR, LG, JL, VT and MC collected the data. SPR compiled, analysed the data and developed the website. SPR and MC wrote the manuscript. All authors reviewed the manuscript.

